# Adaptive landscape of protein variation in human exomes

**DOI:** 10.1101/282152

**Authors:** Ravi Patel, Maxwell D. Sanderford, Tamera R. Lanham, Koichiro Tamura, Alexander Platt, Benjamin S. Glicksberg, Ke Xu, Joel T. Dudley, Laura B. Scheinfeldt, Sudhir Kumar

## Abstract

The human genome contains hundreds of thousands of missense mutations. However, only a handful of these variants are known to be adaptive, which implies that adaptation through protein sequence change is an extremely rare phenomenon in human evolution. Alternatively, existing methods may lack the power to pinpoint adaptive variation. We have developed and applied an Evolutionary Probability Approach (EPA) to discover candidate adaptive polymorphisms (CAPs) through the discordance between allelic evolutionary probabilities and their observed frequencies in human populations. EPA reveals thousands of missense CAPs, which suggest that a large number of previously optimal alleles had experienced a reversal of fortune in the human lineage. We explored non-adaptive mechanisms to explain CAPs, including the effects of demography, mutation rate variability, and negative and positive selective pressures in modern humans. Our analyses suggest that a large proportion of CAP alleles have increased in frequency due to beneficial selection. This conclusion is supported by the facts that a vast majority of adaptive missense variants discovered previously in humans are CAPs, and that hundreds of CAP alleles are protective in genotype-phenotype association data. Our integrated phylogenomic and population genetic EPA approach predicts the existence of thousands of signatures of non-neutral evolution in the human proteome. We expect this collection to be enriched in beneficial variation. EPA approach can be applied to discover candidate adaptive variation in any protein, population, or species for which allele frequency data and reliable multispecies alignments are available.

## Introduction

Over half a million missense variants have been identified in human populations, of which a substantial number occurs at significant frequency (> 1%; 33,369 missense variants) (1000 Genomes Project Consortium 2015). While previous studies have shown the potential for ample adaptive coding variation in the human genome (Boyko et al. 2008; Enard et al. 2014), they have pinpointed only a few missense polymorphisms to be adaptive (Grossman et al. 2013; Hernandez et al. 2011) (**Table 1**). It is possible that virtually all of the common human missense polymorphisms are either selectively neutral or deleterious (i.e., subject to purifying selection), but an alternative explanation is that existing methods lack sufficient power to locate adaptive coding variation. Furthermore, population genomic approaches to date are typically designed to identify recent selective pressures acting on candidate genes or genetic regions that vary within modern human populations, a segment of time that is only a minor fraction of the depth of the human lineage. We, therefore, have the opportunity to discover thousands of novel adaptive changes by using complementary approaches.

**Table 1:** Known adaptive missense polymorphisms and their candidate adaptive polymorphism (CAP) status with empirical probability (*P*neu). *Note*. A candidate adaptive polymorphism (CAP) is an amino acid polymorphism with the evolutionary probability (EP) < 0.05 and population allele frequency (AF) > 5%. n/a marks alleles for which at least one of these two conditions was not met. **Supplementary Table 2** presents more details on each of these polymorphisms and the source references. Marginal status is given to alleles with EP < 0.05 and global AF > 2%.

| Protein | SNP Identifier | CAP? | *P*-value |
| --- | --- | --- | --- |
| ALMS1 | rs10193972 | yes | < 0.02 |
|  | rs2056486 | yes | < 0.02 |
|  | rs3813227 | yes | < 0.02 |
|  | rs6546837 | yes | < 0.02 |
|  | rs6546838 | yes | < 0.02 |
|  | rs6546839 | yes | < 0.02 |
|  | rs6724782 | yes | < 0.02 |
| APOL1 | rs73885319 | no | n/a |
| DARC | rs12075 | yes | < 0.02 |
| EDAR | rs3827760 | yes | < 0.03 |
| G6PD | rs1050828 | marginal | n/a |
|  | rs1050829 | yes | < 0.03 |
| HBB | rs334 | marginal | n/a |
| MC1R | rs1805007 | no | n/a |
|  | rs1805008 | no | n/a |
|  | rs885479 | yes | < 0.03 |
| SLC24A5 | rs1426654 | yes | < 0.02 |
| SLC45A2 | rs16891982 | yes | < 0.02 |
| TLR4 | rs4986790 | yes | < 0.04 |
|  | rs4986791 | marginal | n/a |
| TLR5 | rs5744174 | no | n/a |
| TRPV6 | rs4987657 | yes | < 0.01 |
|  | rs4987667 | yes | < 0.01 |
|  | rs4987682 | yes | < 0.01 |

In this article, we integrate phylogenomics and population genomics to discover candidate adaptive polymorphisms and apply it to the human exome. Our approach advances beyond the current phylogenetic methods that compare patterns across species, but are blind to variation segregating within a given species (Anisimova and Yang 2007; Goldman and Yang 1994; Hurst 2002; Lindblad-Toh et al. 2011; Muse and Gaut 1994; Nielsen et al. 2005; Peter et al. 2012; Pollard et al. 2006; Shapiro and Alm 2008; Yang and Bielawski 2000). It is also distinct from the current population genomic methods that utilize within-population variation to identify candidate adaptive genes or genetic regions, but do not distinguish specific amino acid variants (Akey 2009; Akey et al. 2002; Grossman et al. 2013; Li and Stephan 2006; Moon and Akey 2016; Sabeti et al. 2007; Teshima et al. 2006; Voight et al. 2006). We applied this new approach to over 500,000 polymorphic missense alleles (1000 Genomes Project Consortium 2015) reported in human proteins, which revealed over 18,000 variants that exhibit non-neutral evolutionary patterns. We explored a wide variety of non-adaptive phenomena to explain the existence of these variants and investigated available genotype-phenotype association studies to determine if the non-neutral variants revealed by our new approach have had significant impact on human phenotypic variation.

## New Approaches

Our approach exploits the neutral theory framework, where variation arising from long-term molecular evolution among species informs a null model of observed within-species patterns of selectively neutral variation (i.e., no fitness effect) (Kimura 1983). This relationship is useful to identify adaptive proteins that deviate from neutral expectations and have undergone adaptive evolution (Hudson et al. 1987; McDonald and Kreitman 1991). In our novel allelic approach, we first capture long-term evolutionary history with estimates of the neutral evolutionary probability (EP) of observing each of the possible 20 segregating amino acid residue alleles at a given amino acid position. EP is computed using a Bayesian framework and a multispecies alignment; it is an average of posterior probabilities weighted by the divergence time of each of the species relative to humans in the species timetree used (Liu et al. 2016). The sum of all allelic EPs is 1.0 for each amino acid position. Importantly, EP for an amino acid allele at a given protein position is not affected by the presence of a consensus base at that position in the human reference genome or by the corresponding alleles that segregate in humans, because this information is excluded from the multispecies alignment when EP is calculated (Liu et al. 2016). EP of an allele at a given position is, therefore, completely independent of intra-specific variation. Under neutral theory, residue alleles with low EP (< 0.05) are not expected to persist within populations and are, therefore, predicted to impact function and fitness (Liu et al. 2016). Indeed, less than 1% of simulated neutral EPs fall below 0.05 in computer simulations, where we used the 46 species time tree in **Fig. 1a**, branch lengths from UCSC (Kent et al. 2002; Liu et al. 2016; Murphy et al. 2001; Siepel and Haussler 2005), and pyvolve (Spielman and Wilke 2015) to simulate amino acid sequences (see **Methods**).

**Figure 1:**
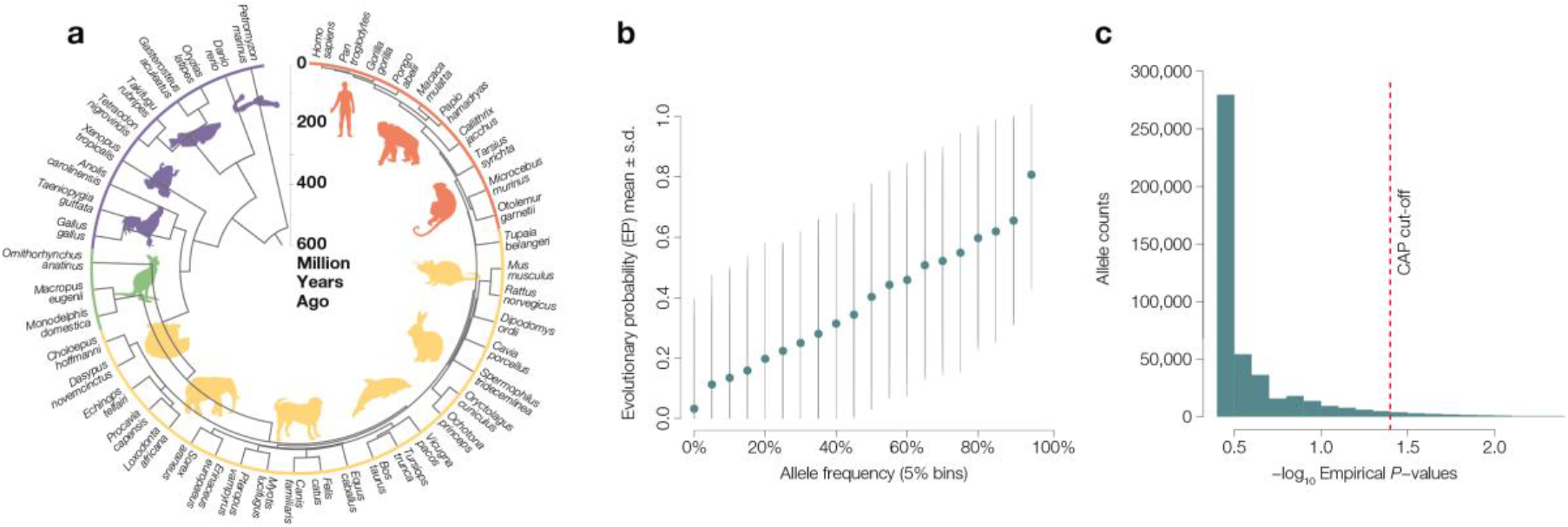
Evolutionary Probability Approach. The evolutionary probabilities (EPs) and their application to discover candidate adaptive polymorphisms (CAPs). **Panel a** displays a timetree of 46 vertebrates and lamprey, including 36 mammalian species, which was used along with alignments of orthologous amino acid sequences for all human proteins(Kent et al. 2002) to compute the probability of observing each amino acid residue at a given position. Under neutral theory, we expect a strong relationship between EP and allele frequency (AF) such that evolutionarily unexpected alleles (EP < 0.05) will be rare. **Panel b** displays the relationship between EP and AF. Average EP (y-axis) was calculated for 0.05 sized AF bins (x-axis) for all polymorphic missense alleles in the 1000 Genomes Project Phase 3 whole genome sequencing data, which confirms the general relationship between EP and AF to be consistent with neutral expectations. The standard deviation is visualized with grey lines (averages are in blue), which is expected to be large because contemporary AFs are a product of time of origin, natural selection, and genetic drift experienced by a mutation. **Panel c** displays the distribution of empirical P values (–log_10_) generated from the empirical framework (AF | EP < 0.05). The cutoff used to identify CAPs is shown with a dashed red line and is more extreme than a false positive rate of 0.05.

Therefore, EP can serve as a null expectation that predicts the neutral probability of observed within-species variation. Contrasting the former against the latter produces a direct neutrality comparison, e.g., non-neutral residue alleles with low EP (< 0.05) are expected to correspond to missense mutations that are found at low allele frequencies (AFs) due to purifying selection (Liu et al. 2016). Consistent with this expectation, 91% of disease-associated missense variants in HumVar (Adzhubei et al. 2010) have low EP (<0.05) and low AF (< 1%). More generally, EP shows agreement with observed global AFs calculated from the 1000 Genomes data (**Fig. 1b**; R^2^ = 0.83, *P* < 10^-15^).

We used the above considerations to build an Evolutionary Probability Approach (EPA) to identify non-neutral (EP < 0.05) alleles that occur with unexpectedly high population AF. When applied to protein sequence variation, such alleles will likely impact protein function, and their prevalence may be due to adaptive pressures. Therefore, we refer to them as candidate adaptive polymorphisms (CAPs). An observed allele is designated a CAP, if it has an EP < 0.05 and AF > 5%. These thresholds were chosen because the empirical probability of observing a CAP for neutral alleles, *P*_neu_, fallsbelow 0.05 for 1000 GenomesProject data (**Fig. 1c**), which represents a significant departure from selective neutrality and forms the basis of EPA. EPA is analogous to empirical outlier approaches frequently utilized in population genomics, including those that identify candidate adaptive polymorphisms with metrics such as *F*_*ST*_ or Tajima’s D (Lewontin and Krakauer 1973; Tajima 1989). A critical difference is that we use information from both phylogenomics (EP) and population genetics (AF) to identify CAPs, which makes EPA a two-dimensional approach and complementary to available methods.

## Results and Discussion

We applied EPA to 515,700 polymorphic missense alleles (1000 Genomes Project Consortium 2015) reported in human proteins. We retrieved EPs for each allele from http://www.mypeg.info (Kumar et al. 2012; Liu and Kumar 2013). The EPs were calculated by Liu et al. (2016) using a 46 species alignment of orthologous amino acid sequences (Kent et al. 2002; Liu et al. 2016). The timetree (Hedges et al. 2006) of these species covers a very large evolutionary timespan (∼5.8 billion years(Hedges et al. 2015); **Fig. 1a**), such that each amino acid position has had ample time to experience mutation and purifying selection.

EPA revealed 18,724 candidate adaptive polymorphisms (EP < 0.05) whose allele frequencies showed significant departure from neutrality (P_neu_ < 0.05). These CAPs were found in 7,815 proteins (see www.mypeg.info/caps for a list of residues) distributed across all autosomal chromosomes (**Fig. 2a**). Many proteins harbor multiple CAPs (**Fig. 3a**), e.g., more than 20 CAPs were found in HLA (**Fig. 2b**) and MUC genes. Both of these gene families play a role in immune response (Parham 2005; Pelaseyed et al. 2014) and are implicated in human adaptation (Andres et al. 2009; Vahdati and Wagner 2016). Several biological processes are significantly enriched for CAP-containing proteins (Mi et al. 2016), including sensory perception, immunity, and metabolism (**Fig. 3b; Supplementary Table 1**).

**Figure 2:**
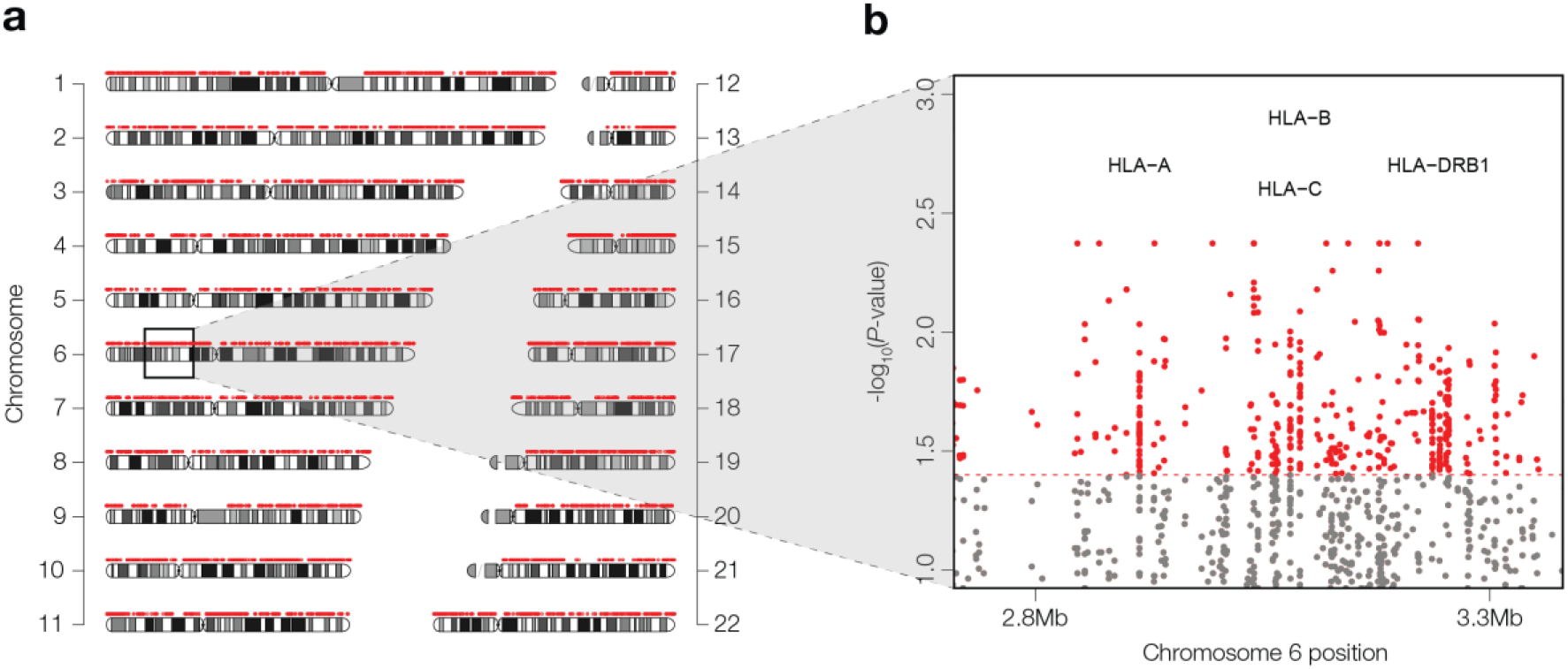
Chromosomal distribution of CAPs.(**a**) The distribution of candidate adaptive alleles (CAPs) across autosomal chromosomes (red points). Chromosomal banding patterns are also visualized for reference. (b) A plot of –log_10_(P_neu_) generated from the Evolutionary Probability Approach (y-axis) against chromosome position (x-axis) for the MHC region of chromosome 6. CAPs are shaded red and non-CAPs are shaded grey. The CAP *P*_neu_ cutoff is shown with a dashed red line. Notable HLA genes with more than 20 CAPs are indicated.

**Figure 3:**
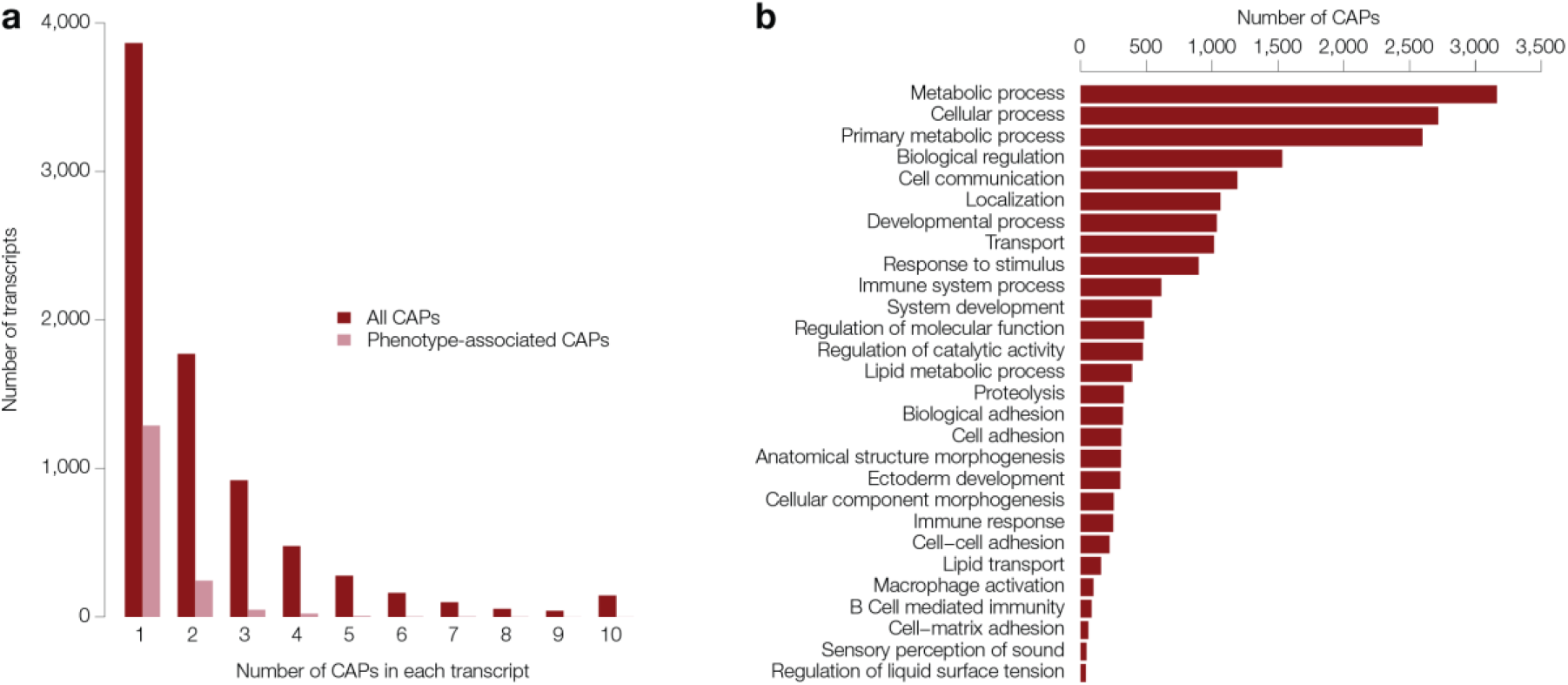
Properties of candidate adaptive alleles. (a) Distribution of all (red bars) and phenotype-associated (pink bars) CAP counts across proteins. (**b**) Biological processes that are significantly enriched for CAPs after Bonferonni correction for multiple testing. The y-axis displays GO-slim biological process category names, and the x-axis displays the number of CAPs annotated to a given GO-slim biological process category. Several categories were significantly enriched with a fold enrichment > 1.5 (**Supplementary Table 1**).

Furthermore, a vast majority (> 70%) of known adaptive amino acid polymorphisms were found to be CAPs (**Table 1; Supplementary Table 2**), which is a significant enrichment (permutation *P <* 10^-7^). EPA also discovers a majority of the protein polymorphisms predicted to be adaptive in previous population genomic analyses (**Supplementary Table 3**), which suggested that the CAP catalog contains many truly adaptive alleles. Still, the size of the CAP catalog is over 200 times larger than the number of previously identified adaptive polymorphisms (**Table 1, Supplementary Tables 2 and 3**).

Previous work would lead us to believe that the majority of common missense mutations are either selectively neutral, in which case allele frequencies are primarily driven by genetic drift, or are mildly deleterious (Kryukov et al. 2007; Zhu et al. 2011), in which case allele frequencies could reflect some combination of drift, compensatory variation, or epistasis. In addition, several non-adaptive phenomena could artificially inflate neutral or deleterious missense allele frequencies. We, therefore, examined the extent to which genomic features and demographic processes could have given rise to CAPs.

### Mutation rate differences and biased gene conversion

Given that mutation rates are known to affect allele frequencies (Harpak et al. 2016), we investigated the potential for mutation rate variation to result in false positive CAPs. We first examined if mutation rates were elevated in codons containing CAPs by comparing the rate of occurrence of synonymous variants in codons that contained CAPs with codons that did not contain CAPs. These two rates were very similar, as 5.7% of the CAP-containing codons also harbored a synonymous polymorphism and 5.4% of non-CAP codons harbored a synonymous polymorphism. This result suggests that mutation rate differences do not explain the observed distribution of CAP allele frequencies.

In addition, the hypermutability of CpG sites did not explain the persistence of low EP alleles at high frequency due to recurrent mutations. We found a smaller proportion of CpG overlapping CAPs relative to non-CAPs (26% and 33%, respectively). Furthermore, we considered whether biased gene conversion could result in false positive CAPs (Ratnakumar et al. 2010). However, fewer than 1% of CAPs were within regions of known biased gene conversion (Capra et al. 2013; Rosenbloom et al. 2015), and the frequencies of weak to strong (W→S) and strong to weak (S→W) changes (Lachance and Tishkoff 2014) for non-CAP alleles (with EP < 0.05 and AF < 5%) were not significantly different than CAP alleles (*P =* 0.90).

### Relaxation of purifying selection

We also examined the possibility that CAP-containing human proteins have experienced relaxation of function in the human lineage. While we think this is unlikely, because it would require a vast fraction of human proteins (> 7,000 out of 22,000) to be under reduced selection, we investigated missense mutations that cause Mendelian diseases and compared the frequency of these mutations in CAP-containing proteins and non-CAP proteins (see **Methods**). We did not find a significant difference in the preponderance of disease mutations in CAP and non-CAP proteins. Therefore, it is unlikely that CAP-containing proteins have become less functionally important relative to other human proteins.

### Adaptive hitchhiking

Deleterious alleles located in genomic regions, which have undergone selective sweeps, can hitchhike to higher than expected frequencies merely due to proximity to and linkage disequilibrium with nearby adaptive alleles (Chun and Fay 2011). Only a small number of CAPs (6.7%) are located in selective sweep regions (Schrider and Kern 2016). This observation is supported by previous studies (Chun and Fay 2011) that investigated the impact of hitchhiking on deleterious allele frequencies and found only a few hundred deleterious hitchhiking nonsynonymous SNPs with common allele frequencies (≥ 5.9%) in the 1000 Genomes Project data. Therefore, hitchhiking of deleterious alleles with selective sweeps does not appear to explain an overwhelming majority of CAPs.

### Human demography

Human demographic history may explain the prevalence of CAPs, because the migration of modern humans out of Africa and subsequent population expansions could have resulted in higher than expected frequencies of deleterious and mildly deleterious alleles. However, it is not likely that these alleles overwhelm the set of CAPs identified, since even a purely neutral model of human evolution does not explain the fraction of alleles found at high allele frequencies: the SFS of empirical CAPs shows a dramatic skew towards high frequency alleles relative to neutral expectation (**Fig. 4a**). We then tested if the CAPs SFS can be generated by human demographic history in combination with various models of selection. We employed a model based on differential equations to approximate the evolution of allele frequencies (Jouganous et al. 2017) and simulated a wide range of negative and positive selection coefficients for a demographic model of recent human history (Gravel et al. 2011) with a range of gamma parameter values (see **Methods**). A model containing negative and positive selections provided the best fit for the CAPs SFS (*lnL* = -3,080; *P* << 10^-10^; **Fig. 4b**). In this model, 47% of the observed alleles were predicted to be weakly deleterious (*s* = -8×10^-4^) and the remaining 53% were beneficial (*s* = +1×10^-3^).

**Figure 4:**
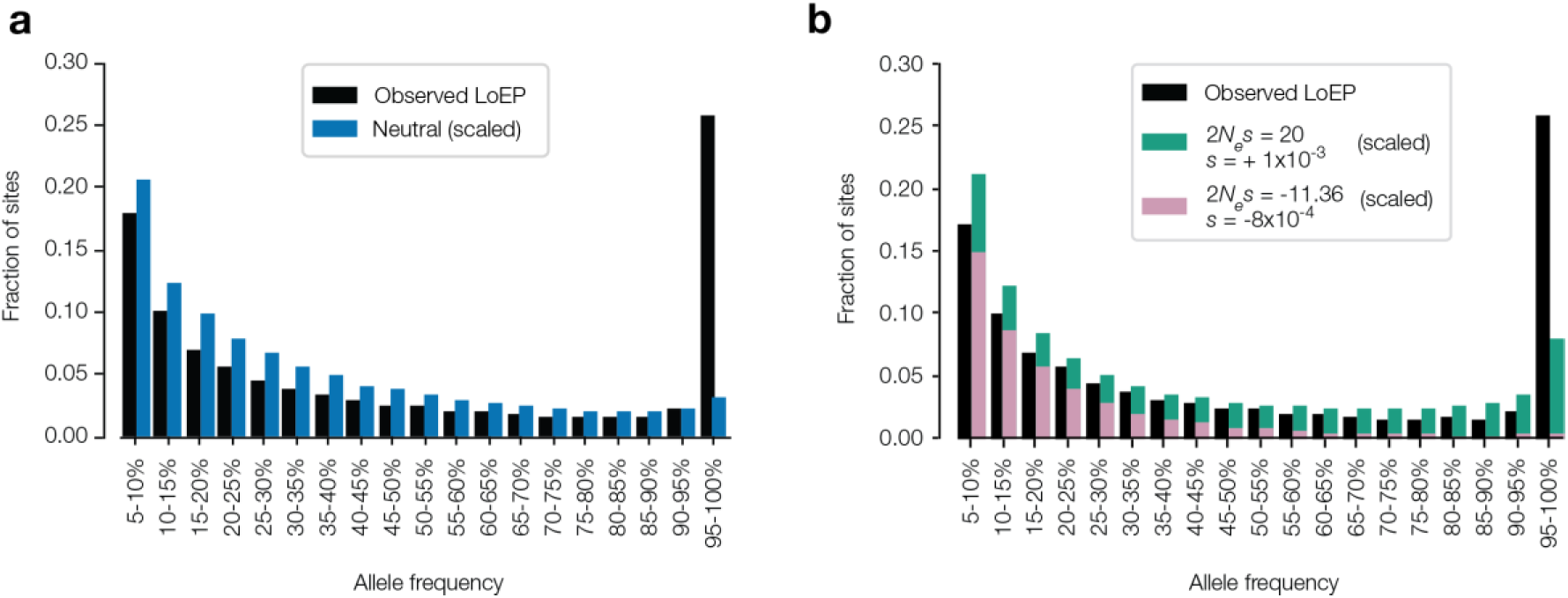
Selection model fits to observed CAPs. Site frequency spectra (SFS) for SNPs with AF > 5%. Site frequency spectra (SFS) were *scaled* to have the same number of sites for AF > 5%. Black bars represent all EP < 0.05 alleles observed in 1000G Phase 3 individuals. (**a**) Observed and fitted SFS for all candidate adaptive polymorphisms (CAPs). A neutral model (blue) does not explain the preponderance of alleles found at very high AF, and does not fit the observed data well (*lnL*= -4,124) (**b**) Observed and fitted SFS for all CAPs. A model with weakly deleterious (purple) and beneficial (green) showed the best fit (*lnL* = -3,080). It was significantly better than any other combination of models (LRT *P* << 10^-10^). All CAP alleles shared with great apes (5%) were excluded from observed SFS.

However, even the best-fit simulated selection model failed to explain the preponderance of polymorphisms with very high frequency (>95%). The number of empirical CAPs in this category was over three times greater than expected (**Fig. 4b**). This result led us to consider whether CAPs were common in the ancestors of modern humans and represent ancestral standing variation. We examined the proportion of CAPs that were shared with archaic hominins (Neanderthals and Denisovans) (Green et al. 2010; Meyer et al. 2012; Prufer et al. 2014) and found that 43% of CAPs are shared with modern humans. This proportion is significantly higher than what is expected by chance (permutation *P* < 10^-7^). While some of the shared CAPs could have resulted from archaic gene flow, the majority of these CAPs were likely present in the last common ancestor of modern humans and archaic hominids, because most (93.6%) shared CAPs occur at very high frequencies (AF > 95%) in modern humans. One such possibility is a CAP (rs4987682) in *TRPV*6, which is present in the Altai Neanderthal genome (Prufer et al. 2014). *TRPV*6 is involved in calcium absorption (Hughes et al. 2008) and located in a region of the genome that has been identified in several previous genome-wide scans for selection (Akey et al. 2006; Hughes et al. 2008). This region is hypothesized to have been subjected to multiple selective events (Hughes et al. 2008).

### Validating CAPs

Generally, traditional functional evaluation of CAPs that arose in the human lineage is challenging, because *in vitro* and *in vivo* approaches are low-throughput, require *a priori* functional information for experimental design, and do not provide the impact of individual alleles on higher-level human phenotypes. Furthermore, it is not possible to test human fitness in a controlled/laboratory setting, and it is often not relevant to test the functional impact of CAPs in non-human model systems. It is, however, possible to take an organismal approach to investigate allelic impact on natural, population-level human variation using phenotype-association studies. For example, many well-known adaptive missense variants (**Table 1**) are also significantly associated with phenotypes in genome-wide studies: rs334 with malaria and severe malaria (Band et al. 2013; Timmann et al. 2012), rs4987667 with intermediate gene expression phenotypes involving HLA (Fehrmann et al. 2011), and rs1426654 with skin pigmentation (Stokowski et al. 2007).

Therefore, we searched the Human Gene Mutation Database (HGMD) (Stenson et al. 2009) for high EP alleles associated with reduced fitness, i.e., the low EP CAP alleles associated with fitness benefits. That is, the evolutionarily preferred allele prior to the divergence of humans and chimpanzees (high EP, EP > 0.5) has experienced a reversal of fortune and become detrimental. We found 253 high EP alleles to be associated with disease phenotypes in contemporary humans, where the low EP CAP allele occurs with AF > 5%.

We also scanned the NHGRI-EBI catalog (MacArthur et al. 2017) of curated GWAS studies to identify additional CAP and found 158 CAPs. Of these, 101 showed odds ratio (OR) less than one for at least one discrete trait related to reduction in the incidence of the associated abnormal phenotype. That is, 60% of the CAPs are protective against the increased disease risk (**Supplementary Table 4**). One such example is a CAP found in the LOXL1 protein that confers a 20-fold decrease in risk for developing exfoliation glaucoma, a leading cause of irreversible blindness (Thorleifsson et al. 2007). Another is *APOE*, which decreases risk five-fold for significant cerebral amyloid deposition (Li et al. 2015). These findings not only suggest functional implications of CAPs, but also that many CAPs are associated with health benefits.

Beyond the limited number of variants in the NHGRI-EBI GWAS catalog, we investigated phenotypic associations in GWAS database that contains a large catalog of genotype-phenotype association studies. We mined data available from GRASP2 (Leslie et al. 2014) to determine whether CAPs have had significant impact on human phenotypes more broadly. We found that 11% of CAPs were significantly associated with tested phenotypes (2,073 alleles at a significance threshold of *P* < 10^-8^), which we refer to as pheno-CAPs. This prevalence of pheno-CAPs is significantly higher than what is expected by chance (permutation *P* < 10^-7^). Moreover, less than 1% of frequency matched non-CAP alleles are significantly phenotype-associated in GRASP2 (*P* < 10^-8^). We tested the possibility that low-EP deleterious recessive alleles have persisted at significant population frequencies. If this had been the case, we would expect an excess of heterozygote CAPs relative to neutral expectations. However, very few CAPs (2.5%) displayed a significant excess of heterozygosity (χ^2^ *P*-value < 0.05). Moreover, after excluding pheno-CAPs that are not shared across all 1000 Genomes continental samples (1000 Genomes Project Consortium 2015), that are located in previously identified selective sweeps (Schrider and Kern 2016), and that are located in previously identified regions containing CpG sites and biased gene conversion regions (Rosenbloom et al. 2015), over 1000 proteins contain one or more pheno-CAPs.

We expect pheno-CAPs to be enriched for causal alleles. There are many reasons for this expectation. First, amino acid polymorphisms alter the sequence of functional genome entities (proteins). Second, if pheno-CAPs are causal alleles then we would expect them to show the strongest association *P*-values among all tested missense variants. This is indeed the case for 92% of CAP proteins, where a pheno-CAP has the strongest association of all missense variants in that protein for a given phenotype in the GRASP2 database (Leslie et al. 2014). Third, a vast majority of putative adaptive variants in humans are CAPs (**Table 1**) and are derived variants in modern-humans; they are not shared with archaic hominins.

In conclusion, we have found over 18,000 missense human polymorphisms that are candidates of beneficial selection. This new adaptive allele catalog is made possible by the EP approach, which is sensitive to a timeframe that predates the out of Africa migration of modern humans, but is not limited to fixed differences between species (Anisimova and Yang 2007; Goldman and Yang 1994; Holt et al. 2008; Hurst 2002; Lindblad-Toh et al. 2011; Muse and Gaut 1994; Nielsen et al. 2005; Peter et al. 2012; Pollard et al. 2006; Shapiro and Alm 2008; Yang and Bielawski 2000). The former timeframe has been addressed by methods that are sensitive to recent classic sweeps and regionally restricted adaptation, which have been the focus of the majority of human adaptation studies to date (Akey 2009; Akey et al. 2002; Grossman et al. 2013; Li and Stephan 2006; Moon and Akey 2016; Sabeti et al. 2007; Teshima et al. 2006; Voight et al. 2006). These studies have yielded only a few adaptive coding variants, leading some to argue that regulatory variation is the predominant raw material for adaptive change (Akey 2009; Fraser 2013; Grossman et al. 2013). Our results suggest that the temporal sensitivity of the EP approach is able to generate a catalog of candidate adaptive polymorphisms that is enriched in functional as well as beneficial variation. We expect many CAPs to be involved in compensatory evolution and synergistic epistasis to counter genetic load exerted by deleterious variants that have risen to high frequencies due to human demography and genetic drift. Therefore, CAPs provide ready hypotheses to test in future computational and experimental investigations.

## Materials and Methods

### 1000 Genomes Allele Frequencies

Global allele frequencies (AFs) for all missense single nucleotide polymorphisms (SNPs) (*n* = 515,700) in the 1000 Genomes Project phase 3 data (1000 Genomes Project Consortium 2015) were calculated for all unrelated individuals (*n* = 2,405). More specifically, one of each related pair of individuals identified in the Phase 3 release (ftp://ftp.1000genomes.ebi.ac.uk/vol1/ftp/release/20130502/20140625_related_individuals.txt) was removed before calculating global allele frequencies. For each polymorphic nucleotide position, EP estimates for the codons corresponding to the reference (hg19) and non-reference nucleotides were used. For each allele, we tested for an overrepresentation of potentially deleterious recessive CAP heterozygotes and evaluated the proportion of CAPs that were in Hardy-Weinberg (HW) disequilibrium (HW χ^2^ *P*-value < 0.05).

### Evolutionary Probabilities

Evolutionary probabilities (EPs) were calculated for each amino acid residue using the method of Liu et al.(Liu et al. 2016) and a 46 species alignment of orthologous amino acid sequences(Kent et al. 2002; Liu et al. 2016) (they are available from http://www.mypeg.info (Kumar et al. 2012; Liu and Kumar 2013)). The timetree (Hedges et al. 2006) of these species covers a very large evolutionary timespan (∼5.8 billion years (Hedges et al. 2015); **Fig. 1a**), such that each amino acid position has had ample time to experience mutation and purifying selection. We designed a simulation to verify that the EP was over 0.05 for neutral alleles, by using the 46 species time tree in **Fig. 1a** and branch lengths from UCSC(Kent et al. 2002; Liu et al. 2016; Murphy et al. 2001; Siepel and Haussler 2005). Using pyvolve v0.8.7 (Spielman and Wilke 2015), we generated 1000 replicate datasets of proteins with 500 amino acid positions and calculated EP for alleles at each site.

### Evolutionary Probability Approach Framework

We began with the premise that for a given amino acid position, the probability the position has been neutral (EP) over long-term evolutionary history (inferred from inter-species comparisons as described in (Liu et al. 2016)) combined with the orthogonal shorter-term intra-specific purifying and directional selective pressures (captured by population allele frequency, AF) produces a categorical framework for genome-wide variation. This framework distinguishes neutral, potentially deleterious, and potentially adaptive variation. The sum of all allelic EPs is 1 for each amino acid position, and residues with low EP (< 0.05) are unexpected under neutral theory (Liu et al. 2016). We developed an empirical framework to identify candidate adaptive polymorphisms (CAPs): Prob(AF | EP < 0.05), and for each allele, calculated a one-sided cumulative empirical *P*-value using a cumulative distribution function (CDF) implemented with a custom R script (R Core Team 2014).

### Misinference of ancestral state

In genomic scans for selection, misidentification of ancestral states may cause false signatures of selection (Baudry and Depaulis 2003). EPA fortunately does not suffer from this problem, because it requires EP < 0.05. An allele with such a low EP will likely arise in the human lineage after their divergence from chimpanzees. Additionally, EP calculation utilizes a probabilistic model that integrates over all the outgroup species in an alignment, which makes it better than methods that utilize one or a few outgroups to properly identify the derived allele (Hernandez et al. 2007; Keightley et al. 2016). Consistent with this property, we did not find any CAP alleles in all three of the Great Ape species (chimpanzee, gorilla, and orangutan) in our multispecies protein alignments. A comparison with chimpanzee proteins revealed 3.5% CAP allele sharing, and gorilla and orangutan showed 0.7% and 1.1% CAP allele sharing, respectively, with humans. We excluded all of these alleles from all the population genetic analyses, because these CAP residues may have arisen prior to the origin of human lineage.

### Identifying allele sharing with archaic genomes

To determine allele sharing among modern humans and archaic hominins, we collected genome sequencing data for five archaic hominins (four Neanderthal individuals, and one Denisovan individual). One Neanderthal sequence and one Denisovan sequence were acquired from the Max Planck Institute for Evolutionary Anthropology site (http://cdna.eva.mpg.de/neandertal/altai/Denisovan). The three remaining Neanderthal alignments were retrieved from the UCSC Neanderthal Sequence Track (https://genome.ucsc.edu/cgi-bin/hgTrackUi?db=hg19&g=ntSeqReads). We only used sequences that provided > 45% genomic coverage. We defined an allele as shared if it was present in any of these five archaic individuals. A shared allele can be polymorphic or fixed in this aggregated archaic sample.

### Scanning Genotype-Phenotype Association Catalogs

We scanned 75,810 phenotype associated missense mutations in the Human Gene Mutation Database (HGMD) (Stenson et al. 2009) for those that occur at CAP sites. We found 973 such mutations, which we checked for high EP risk-alleles (causing the abnormal phenotype). A high EP risk allele at a CAP site was considered a “reversal”, since this previously favored allele (based on EP) leads to an unfavorable phenotype. We also scanned the NHGRI-EBI GWAS catalog (MacArthur et al. 2017) (January 16, 2018 update) for similar reversals. Filtering the SNPs, we find 158 missense mutations at CAP sites. The NHGRI-EBI GWAS Catalog always reports the risk-allele (the allele that increases phenotypic measurement, e.g., increases disease risk). In order to determine the odds ratio (OR) for the CAP allele, which is often not the reported risk allele, we calculated the inverse (1 / reported OR) when the risk allele was in fact the reversal (high EP allele). An OR < 1 indicates that the allele confers a decrease in abnormal phenotype risk, while an OR > 1 indicates that the allele increases risk for the associated abnormal or case phenotype. Multiple associations were occasionally found for CAPs in the GWAS catalog. We simply reported the study that had the lowest risk-factor (OR) for abnormal phenotypes per CAP allele found.

### Gene Ontology Enrichment

We used the Panther Classification System (Mi et al. 2016) to test for enrichment of Gene Ontology (GO slim) biological processes. As input, we used the list of protein IDs that contain one or more CAPs. We excluded terms with less than two proteins, and we adjusted enrichment *P* values to account for multiple testing with a Bonferroni correction.

### Demographic Simulations

We performed 10,000 forward simulations of human history for 58,000 generations before current time; the simulation scheme includes the out-of-Africa migration of humans (OoA), as well as a subsequent split between simulated European and East Asian populations. The population model includes three representative continental groups (African, European, East Asian). SLiM2 (Haller and Messer 2017) was used for the simulations, with parameters obtained by Gravel et. Al (Gravel et al. 2011). Using a modified SliM2 script to output MS (Hudson) format chromosomes, we sampled individual sequences (50,000 base pairs in length) from the simulated populations at each of the following time points:

(a) the generation immediately before the OoA split (ancestral population), (b) the generation immediately before the European and East Asian split, (c) the contemporary African population, (d) the contemporary European population, and (e) the contemporary East Asian population. Using allele frequencies (AF) from these samples, we followed variants at different AF (0.1%, 1%, and 10%) in the ancestral population and traced their trajectories into the modern day human populations (contemporary populations). For each of these variants, we determined the fraction that achieved > 5% AF (required for CAP status), and were shared among one, two, and three of the contemporary population samples.

### Simulating selection and fitting distributions of fitness effects

We simulated site frequency spectra (SFS) using Moments (Jouganous et al. 2017) to infer distributions of fitness effects (DFE) that explain CAPs for which the human alleles were not shared with any of the three great ape species (chimpanzee, gorilla, and orangutan). Using *dadi* (Gutenkunst et al. 2009), we calculated multinomial log-likelihoods (*lnL*s) of the observed data (CAPs) for simulated deleterious, neutral, and beneficial selection models (as above). We also calculated *lnL* of DFE fit for all possible combinations: deleterious and neutral; neutral and positive; deleterious and beneficial; and, deleterious and, neutral, and beneficial. In this case, we used a single point mass fixed for each type of selection and explored various 2N_e_s values. The model with the highest lnL provides thebest fit for the observed data. We excluded all CAPs shared with great apes in these analyses. The best fit model and *lnL* values for all the CAPs are shown in **Fig. 4b**. We used likelihood fits and Akaike information criterion (AIC) to select the best model.

### Examination of the Relaxation of purifying selection

We examined the possibility that CAP-containing human proteins have experienced relaxation of function in the human lineage. We investigated missense mutations that cause Mendelian diseases and compared the frequency of these mutations in CAP-containing proteins and non-CAP proteins. This analysis used the HumVar (Adzhubei et al. 2010) dataset and obtained the number of disease mutations normalized by the total sequence length and evolutionary rate of CAP and non-CAP proteins. This normalization is required because longer proteins are known to contain more disease mutations as do slower evolving proteins (Miller and Kumar 2001). The ratio of two normalized counts was 0.98, which is close to the expected value of 1.0 corresponding to no difference in the preponderance of disease mutations in CAP and non-CAP proteins.

### Permutation Testing

In order to determine whether the observed proportion of CAPs that have been previously identified as adaptive in humans is higher than would be expected by chance, we randomly sampled 18,724 variants from the set of all human missense variants (regardless of EP), and calculated N_sim_, which captures how often the simulated proportion of phenotype-associated variants was as high or higher than the empirical result. In total, we ran 10^6^ permutations, and calculated a permutation *P-*value with the following equation: (N_sim_ + 1)/1000001.

Similarly, we tested whether the observed proportion of CAPs that are shared with archaic genomes is higher than would be expected by chance. We randomly sampled 18,724 variants from the set of all human missense variants, andcalculated N_sim_, which captures how often the simulated proportion of archaic-shared variants was as high or higher than the empirical result (6,916 for *P* < 0.05 and 2,075 for *P* < 10^-8^). In total, we ran 10^6^ permutations, and calculated a permutation P-value with the following equation: (N_sim_ + 1)/1000001.

In order to determine whether the observed proportion of CAPs that are also associated with phenotypes in the GRASP2 database (Leslie et al. 2014) is higher than would be expected by chance, we randomly sampled 18,724 variants from the set of all human missense variants with an AF > 1% (regardless of EP), and calculated N_sim_, which captures how often the simulated proportion of phenotype-associated variants was as high or higher than the empirical result (6,916 for *P* < 0.05 and 2075 for *P* < 10^-8^). In total, we ran 10^6^ permutations, and calculated a permutation P-value with the following equation: (N_sim_ + 1)/1000001.

## Acknowledgements

We thank Drs. Jody Hey, Rob Kulathinal, Joshua Shraiber, and Heather Rowe for their critical comments on previous versions of this manuscript. We would also like to thank Michael Li and Keith Davis for technical assistance. This work was funded by research grants from NIH (R01HG008146-01 and R01DK098242-04).

## Author Contributions

S.K., L.B.S., and R.P. designed the research study, directed the analysis, and wrote the manuscript, A.P. designed one analysis, and contributed to the manuscript, T.R.L. conducted analyses, M.S. helped with data collection, web development, and analysis, and K.T., B.S.G., K.X., and J.T.D assisted with statistical analysis and contributed to the manuscript.

